# Small teams of myosin Vc coordinate their stepping for efficient cargo transport on actin bundles

**DOI:** 10.1101/106658

**Authors:** Elena B. Krementsova, Ken’ya Furuta, Kazuhiro Oiwa, Kathleen M. Trybus, M. Yusuf Ali

## Abstract

Myosin Vc (myoVc) is unique among vertebrate class V myosin isoforms in that it requires teams of motors to transport cargo. Single molecules of myoVc cannot take multiple steps on single actin filaments, in stark contrast to the well-studied myosin Va (myoVa) isoform. Consistent with *in vivo* studies (1), only teams of myoVc motors can move continuously on actin bundles at physiologic ionic strength (2), raising the question of how motor motor interactions cause this preference. Here, using DNA nanostructures as synthetic cargos for linking defined numbers of myoVa or myoVc molecules, we compared the stepping behavior of myoVa versus myoVc teams, and myoVc stepping patterns on single actin filaments versus actin bundles. Run lengths of both myoVa and myoVc teams increased with motor number, but the run lengths of myoVc teams were longer on actin bundles than on filaments. By resolving the stepping behavior of individual myoVc motors with a Qdot bound to the motor domain, we found that coupling of two myoVc molecules significantly decreases futile back/side steps, which were frequently observed for single myoVc motors. Data showing how changes in the inter-motor distance of two coupled myoVc motors affected stepping dynamics suggested that mechanical tension coordinates the stepping behavior of two molecules for efficient directional motion. Our study thus provides a molecular basis to explain how teams of myoVc motors are suited to transport cargos such as zymogen granules on actin bundles.

**Abbreviations:**

MyoVc: Myosin Vc
myoVa: Myosin Va
Qdots: Quantum dots
SD: Standard Deviation
SE: Standard Error
DNA: deoxyribonucleic acid

## Introduction

In vertebrates, three class V myosin isoforms transport intracellular cargoes such as melanosomes, secretory vesicles, endoplasmic reticulum, and mRNA on actin tracks. MyoVa is the most well-characterized isoform, and the prototype of a processive myosin, i.e. a single motor that can take many steps on an actin filament without dissociating. More limited assays with myoVb suggested that it is also processive (3). In contrast, myosin Vc, which is predominantly expressed in glandular tissues such as pancreas, colon and stomach, was the only vertebrate class V isoform to be characterized as non-processive, implying that it would be unable to move cargo continuously as a single motor (2-4). This observation was surprising because myoVc is known to be involved in transport of secretory vesicles to the apical membrane (5). It was recently shown that in the exocrine pancreas, parallel bundles of actin filaments nucleated by formins at the plasma membrane are the tracks on which zymogen granules are trafficked (1). Consistent with this biological observation, we recently showed that actin bundles are the required track for ensembles of myoVc to continuously move a cargo at physiologic ionic strength (150 mM KCl). Single myoVc molecules can also walk processively on actin bundles, but only at low ionic strength (25 mM KCl) (2). The preference of myoVc for actin bundles is because myoVc takes ~40% back steps and lateral steps under unloaded conditions, and has a very broad turning angle distribution compared with the highly processive myoVa motor (2), suggesting that only the additional binding sites provided by an actin bundle allows for continuous motion. Chimeric constructs identified the lever arm/rod domains of myoVc as the structural elements responsible for this unusual stepping behavior, because the myoVc motor domain fused to the lever arm/rod of myoVa exhibited no back steps and moved processively on single actin filaments, similar to myoVa (2). Although the lever arm/rod junction of myoVa is flexible (Michalek 2015), this isoform moves processively on a single actin filament.

The requirement for teams of myoVc to move continuously on actin bundles near physiologic ionic strength prompted us to further investigate the ensemble behavior of myoVc. Sakamoto and colleagues (6) showed for the first time that when two myoVc motors were coupled via a DNA scaffold, the complex moved continuously at low ionic strength even on single actin filaments. Two motors increased the probability of at least one myoVc head remaining bound to the filament at any given time. Numerous studies have shown that intracellular cargos are transported by the coordination of multiple motors. Melanosomes, for example, are associated with as many as 60 myoVa motors *in vivo*, although it is likely that only a subset of this number are actively engaged with the track at any given time (7). The mechanism by which the mechanical properties of individual motors contribute to or share their collective transport duties depends on the particular motor or motors involved (7).

A number of studies have predicted the collective behavior of motor ensembles transporting a common cargo, but the number of motors and their stepping pattern were generally unknown (8-10). In previous studies with myoVc ensembles (2,6), the stepping dynamics of individual heads in the ensemble were not followed. Here, we designed DNA origamis or DNA scaffolds to act as a synthetic cargo for linking defined number of Qdot-labeled myoVc motors with defined inter-motor spacing (11,12). This approach allowed us to follow the stepping dynamics of two myoVc motors on actin filaments and actin bundles, and to compare this behavior with myoVa. We confirmed that a single molecule of myoVc walks on an actin bundle with frequent back and side steps, but further showed that coupling two myoVc motors significantly decreases the frequency of such futile steps, and causes both the leading and trailing motors to coordinate. Run length enhancement of myoVc by multiple myoVc motors is significantly higher on actin bundles than on actin filaments, in contrast to multiple myoVa motors which show a greater run length enhancement with motor number on single actin filaments. This study provides a molecular basis to explain how multiple myoVc motors are optimized to move large exocrine secretory granules on actin bundles in the final stages of secretion(1).

## Results

### A single myoVc motor on actin filaments or actin bundles

Here we used DNA nanostructures as synthetic cargos for linking defined numbers of myosin molecules. To confirm that the attachment of the DNA origami does not affect the motion of myoVc, a single, dimeric myoVc motor was attached to a Cy3 conjugated DNA nanotube via a SNAP-ligand binding site (see Experimental Procedures). The behavior of motor-origami complexes were observed on single actin filaments that were immobilized onto the glass surface using a Total Internal Reflectance Fluorescence (TIRF) microscope with high temporal (33 ms) and spatial resolution (6 nm). Consistent with previous observations (2,6), single molecules of myoVc did not move processively on actin filaments. In a complementary experiment, the motor domain of a single myoVc motor was labeled with a Qdot (see Experimental Procedures), and again no motion was observed. These results suggest that the attachment of the DNA origami does not affect the behavior of myoVc. We next confirmed the recent observation (2) that single myoVc motors can take multiple steps on actin bundles at 25 mM KCl (Fig. 1A,B). We observed that the velocity obtained using a Qdot bound to the motor domain of myoVc (128 ±41 nm/s. mean ± SD, N=81) was not significantly different from that obtained using the motor-origami complex (116 ± 38 nm/s, N=85), again suggesting that the attachment of the DNA origami does not affect the motion of myoVc. The mean run length of myoVc on actin bundles was determined from an exponential decay distribution function, and velocity was measured from the fit to a Gaussian distribution function. The average run length of a single myoVc molecule on actin bundles was 0.40 ± 0.07 μm (mean ± SE, N=85), and the velocity was 116 ± 38 nm/s (mean ± SD, N=85) (Fig. 2A,C).

**FIGURE 1.**
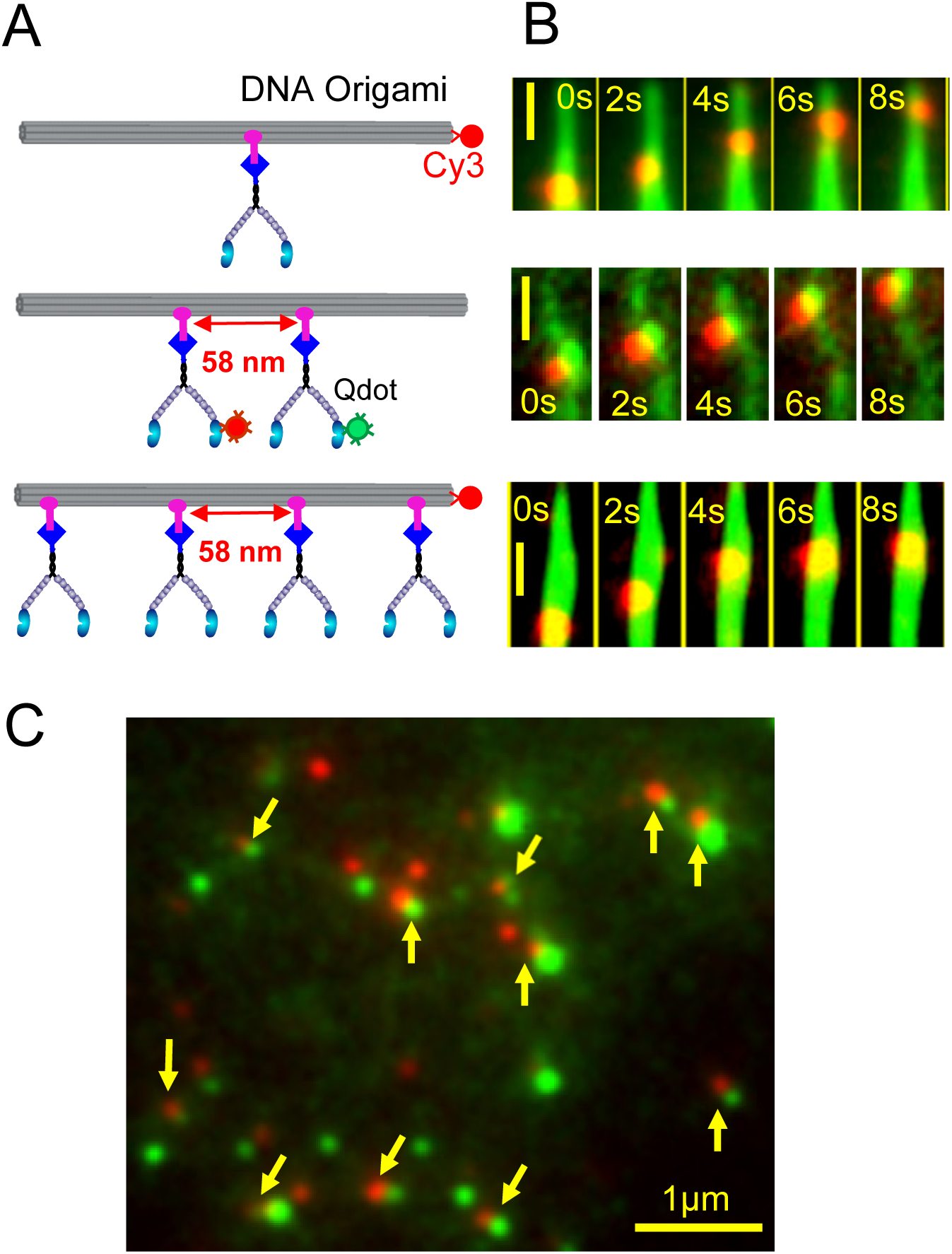
Experimental design to visualize motility of small ensembles of myosin. (A) 1, 2, or 4 myosin motors containing a SNAP-tag at the C-terminus of the myosin heavy chain (blue diamond) are bound to a DNA origami containing varying numbers of SNAP-ligands. Single and 4-motor complexes were visualized using Cy3 (red circle) attached to the DNA origami. In the 2-motor complex, one motor domain of each of the two molecules was labeled with a different colored Qdots (red or green). (B) (Top) Representative sequential images of a single motor (red) moving on an Alexa 647 a labeled actin bundle (green), (middle) a 2-motor complex (red and green dots) moving on an unlabeled actin filament, (bottom) a 4-motor complex (red) moving along an Alexa 647 (green) labeled actin bundle at 1 mM MgATP. Scale bar, 500 nm. Elapsed times are indicated. (C) Data showing that two different colored motors (red and green, yellow arrow) can be coupled to the 2-motor DNA origami. Only the motility of dual colored images were quantified. Scale bar, 1μm.

**FIGURE 2.**
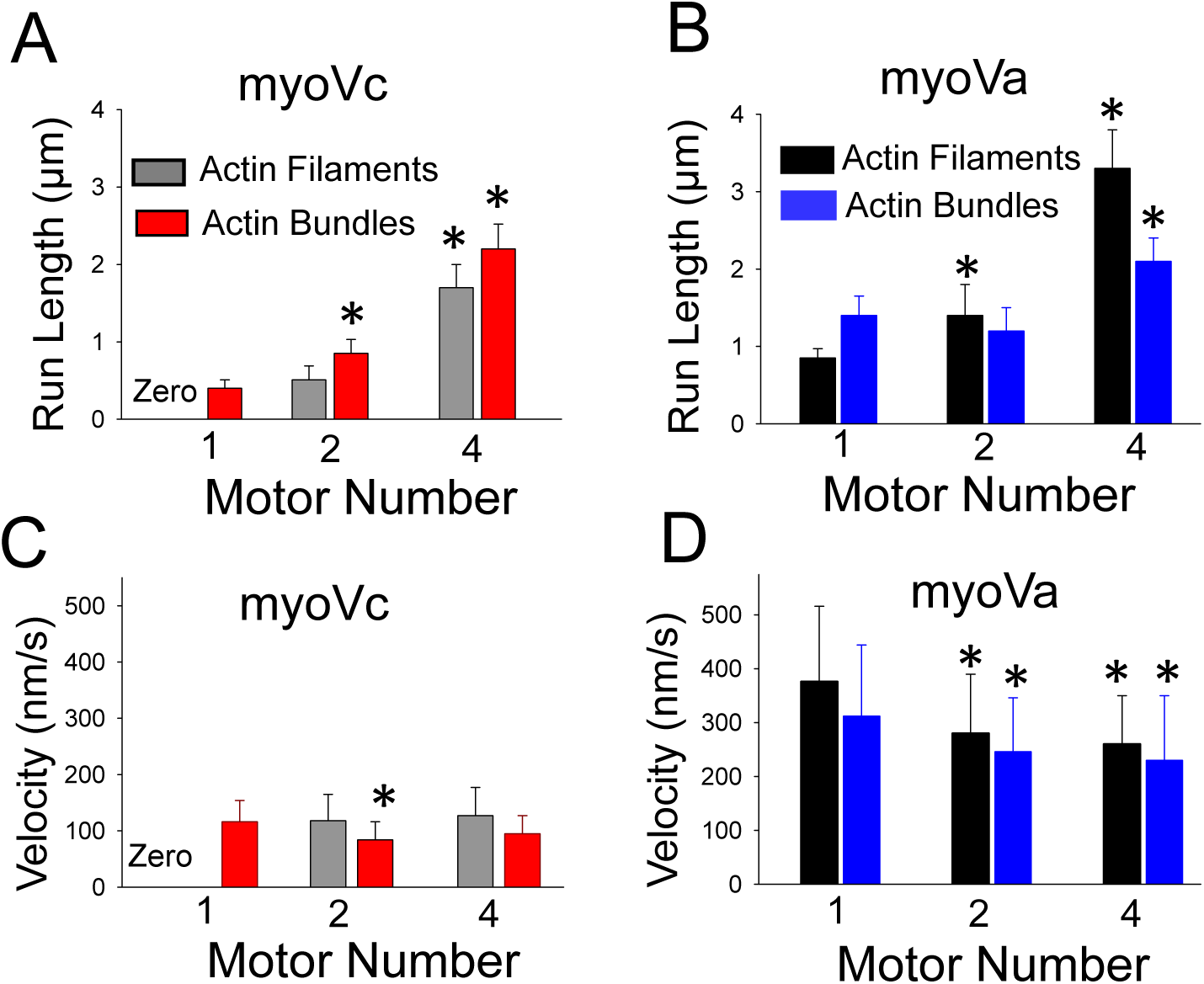
Effect of motor number on run length and velocity of myoVc versus myoVa on actin filaments and bundles. (A) MyoVc run length increased significantly (*, p<0.05) with motor number on both single actin filaments (gray) and actin bundles (red). The run length of a single myoVc on actin bundles is 0.4 ± 0.07 μm (N=85), 0.85 ± 0.14 μm (N=62) for the 2-myoVc complex, and 2.2 ± 0.32 μm (N=48) for the 4-myoVc complex (red). On single actin filaments, the run length of 2 and 4-myoVc complexes are 0.51 ± 0.11 μm (N=54) and 1.7 ± 0.3 μm (N=50) (gray), respectively. The run length at any given motor number is significantly higher (*, p<0.05) on actin bundles than on single actin filaments. The 2-myoVc run length on actin bundles (0.85 ± 0.14 μm, N=62) was 66% longer (p<0.05) than on single actin filaments (0.51 ± 0.11 μm, N=54). The 4-myoVc motor complex run length on actin bundles (2.2 ± 0.32 μm, N=48) is 29% higher than on actin filaments (1.7 ± 0.3μm, N=50). Values in panel A are mean ± SE. Conditions: 25 mM imidazole, pH 7.4, 4 mM MgCl_2_, 1 mM EGTA, 25 mM KCl, 10 mM DTT. (B) The run length of myoVa increased significantly (p<0.05) as a function of motor number on single actin filaments (black). The run length of the 2- (1.4 μm ± 0.4 μm, N=48) and 4-myoVa complex (3.3 μm ± 0.49 μm, N=47) on actin filaments was 1.6 and 3.9 fold longer, respectively, than that of a single myoVa (0.85 ± 0.12 μm, N=65). On actin bundles (blue), the run length of myoVa was decreased for the 2-myoVa complex (1.2 ± 0.3 μm, N=50) but increased significantly for the 4-myoVa complex (2.1 ± 0.3 μm, N=60) compared with a single motor (1.4 μm ± 0.25 μm, mean ± SE; N=58) (Fig. 2B, blue). Values in panel B are mean ± SE. Conditions: 25 mM imidazole, pH 7.4, 4 mM MgCl_2_, 1 mM EGTA, 25 mM KCl, 10 mM DTT. (C) MyoVc velocities were not statistically different except for the 2-myoVc complex on actin bundles (red) (p<0.05) compared with a single myoVc. The velocity of the 2-myoVc complex (84 ± 32 nm/s, N=62) was reduced by 28% compared (p<0.05) with a single motor (116±38 nm/s, N=85), but the velocity of the 4-myoVc complex (95 ± 32 nm/s, N=55) was the same as a single motor. On single actin filaments (gray), the velocity of the 2- (118 ± 47 μm/s, N=60) and 4-myoVc (127 ± 50 nm/s, N=55) complex were the same. Values in panel C are mean ± SD. Conditions: 25 mM imidazole, pH 7.4, 4 mM MgCl_2_, 1mM EGTA, 25 mM KCl, 10 mM DTT. (D) The velocity of 2- or 4-myoVa motor complexes were reduced significantly (p<0.05) compared with a single motor on both single actin filaments (gray) and bundles (blue). Velocity for the 2 versus 4-myoVa complex was not significantly different on either single actin filaments or actin bundles. On single actin filament, the velocity of a single, 2- and 4-myoVa complexes are 376 ± 96 nm/s (N=70), 280 ±110 nm/s (N=68), and 260 ± 90 nm/s (N=60), respectively. On actin bundles, these values are 312 ±132 nm/s (N=68), 246 ± 101 nm/s (N=64), and 230 ± 120 nm/s (N=54), respectively. Values in panel D are mean ± SD. Conditions: 25 mM imidazole, pH 7.4, 4 mM MgCl_2_, 1 mM EGTA, 25 mM KCl, 10 mM DTT.

### Comparison of movement by small ensembles of myoVc versus myoVa

The motion of small ensembles of myoVc or myoVa was observed by coupling motors to a DNA origami containing two or four SNAP ligands spaced at 58 nm intervals (Fig. 1A). An advantage of the DNA origami is that its rigidity keeps the average attachment spacing between the motors constant during movement. Motion of motors was tracked by Qdots attached to the motor domain in the case of the 2-motor complex, or by Cy3 attached to the DNA origami for the 4 motor complex (Fig. 1A).

In contrast to a single myoVc motor, two or four myoVc motors bound to a DNA nanotube moved continuously on single actin filaments at 1 mM MgATP and 25 mM KCl (Fig. 1B), where the ionic strength is considerably lower than physiologic (150 mM KCl). This result was consistent with a previous observation showing that two myoVc motors coupled to a DNA scaffold move continuously at low ionic strength on single actin filaments (6). In our case, the 2-motor complex was formed from equimolar amounts of red and green Qdot labeled myoVc, which enabled tracking of one head of both motors attached to the DNA origami during motion. We observed that 40% (N=165) of complexes were labeled with dual color Qdots (Fig. 1C). Given that statistically a maximum of 50% of the DNA origamis could be dual color (red and green), the binding affinity of myoVc motors to the origami and labeling with Qdots must both be very efficient (80%) (Fig. 1C; see Experimental Procedures). For the 4-motor complex, a five-fold molar excess of motor over SNAP-ligand on the origami was used to favor full occupancy.

On single actin filaments, the run length of both myoVc (gray bars) and myoVa (black bars) increased with motor number (Fig. 2 A,B). On actin bundles, the run length of 2- and 4-myoVc complexes were 112% and 450% longer than that of a single motor (Fig. 2A, red bars). In contrast, the run length of myoVa was decreased by 14% for the 2-motor complex and increased only 50% for the 4- motor complex compared with a single motor (Fig. 2B, blue bars). Comparing motor performance on single actin filaments versus bundles, myoVc and myoVa also differed. For both 2- and 4-myoVc complexes, the run length on bundles was statistically longer than on single actin filaments (Fig. 2A). In contrast, the run lengths of 2- and 4- myoVa complexes on actin bundles were 14% and 36% shorter than that on single actin filaments (13) (Fig. 2B). These results are consistent with the idea that myoVc is specialized for cargo transport on actin cables (2).

For myoVc, velocity of 2- and 4- motor complexes were not staistically different on actin filaments or actin bundles, and velocities on actin bundles were either the same or only slightly less than on filaments (Fig. 2C). However, the velocity of the 2-myoVc complex (84 ± 32 nm/s, N=62) was reduced by 28% compared (p<0.05) with a single motor (116±38 nm/s, N=85). In contrast, velocities of myoVa ensembles were statistically slower than single motors, regardless of whether the motor moved on single actin filaments or bundles (Fig. 2D). This reduction in speed is similar to that observed in previous studies in which two myoVa motors were linked via a DNA scaffold or a Qdot (13,14).

### MyoVc movement at physiological ionic strength

Our previous results with multiple myoVc motors attached to a Qdot showed that even teams of motors required actin bundles for continuous motion at physiologic ionic strength (2). Consistently, the multi-motor myoVc complexes coupled to the DNA origami could not move continuously on single actin filaments at 150 mM KCl, but showed continuous motion on actin bundles, with a run length of 0.39 ± 0.06 μm (mean ± SE, N=64) for the 2-myoVc complex, and 1.4 ± 0.3 μm (mean ± SE, N=58) for the 4-myoVc complex (Fig. 3A). This result implies that both multiple myoVc motors and actin bundles are required for transporting secretory vesicles in the cell (1,5). The run length of 2- and 4-motor complexes was significantly lower at 150 mM KCl compared with what was observed at 25 mM KCl (Fig. 2A versus 3A). At 150 mM KCl, the velocity of the 2-motor complex (120 ± 47 nm/s, N=64) was 43% higher (p<0.05) than that at 25 mM KCl (84 ±32, mean ± SD, N=62), but the velocity of the 4- motor complex was not significantly different at the two salt concentrations (109 ± 46 nm/s, N=58 at 150 mM KCl; 95 ± 32 nm/s, N=55 at 25 mM KCl; mean ± SD) (Fig. 2C versus 3B).

**FIGURE 3.**
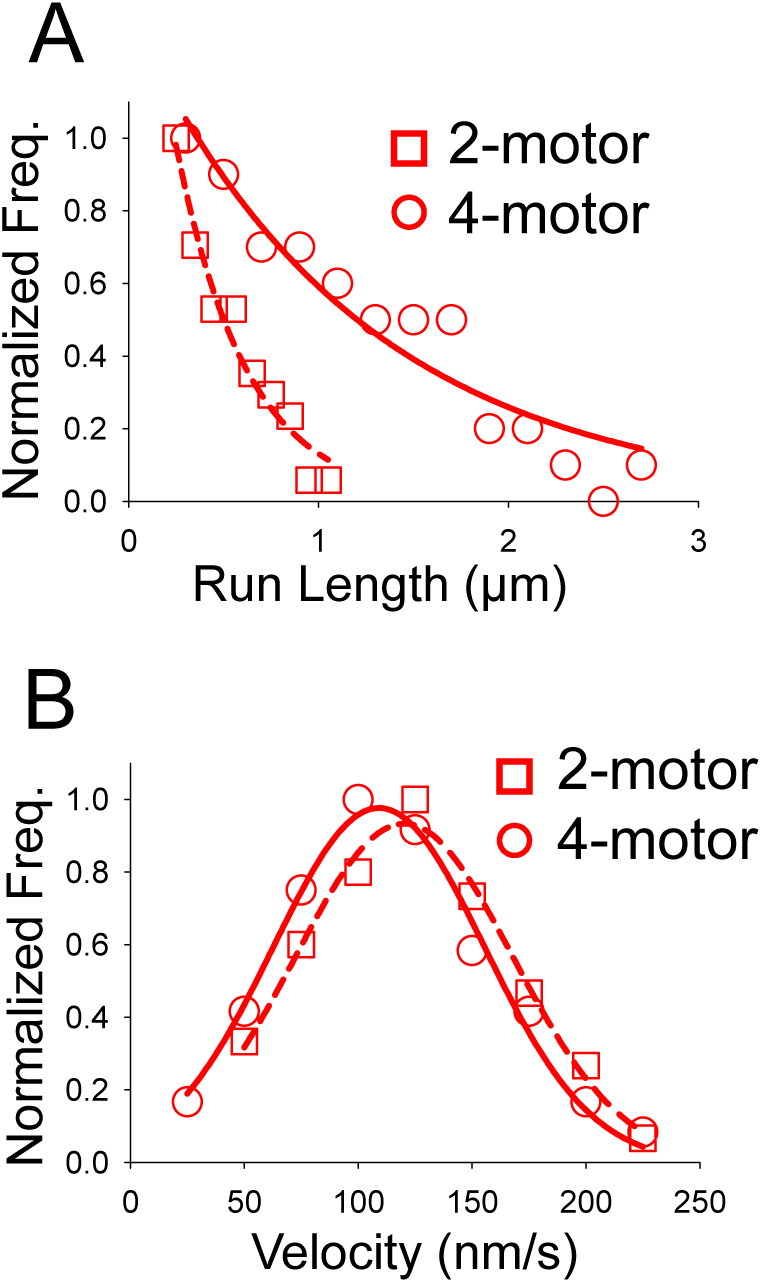
Run length and velocity of myoVc at physiological salt concentration. (A) At 150 mM KCl, the run length of a 4-myoVc complex (1.4 ± 0.3 μm, N=58) on actin bundles was 3.5-fold longer than that of a 2-myoVc complex (0.39 ± 0.06 μm, N=64). Values are mean ± SE. (B) Velocity of the 2-myoVc complex on actin bundles is not significantly different from that of a 4-myoVc complex (120 ± 47 nm/s, N=64 versus 109 ± 46 nm/s, N=58). Values are mean ± SD. (Conditions: 25 mM imidazole, pH 7.4, 4 mM MgCl_2_, 1 mM EGTA, 150 mM KCl, 10 mM DTT).

### Inter-motor distance and cargo rigidity affect run length

We designed three different DNA scaffolds containing two binding sites separated by either 22 nm, 39 nm or 69 nm (Fig. 4A). The DNA scaffold was formed from double-stranded DNA and is less stiff than the DNA origami which was constructed from 6 single DNA strands. The run lengths of the 2-motor complex with 36 nm or 69 nm motor spacing on single filaments were significantly higher than that of the complex with 22 nm spacing, suggesting that spacing between motors is an important factor for efficient cargo transport (Fig. 4B). The shorter run length at 22 nm spacing might be the consequence of steric interference when the binding site for the trailing motor is already occupied by the leading motor. In such a situation, both trailing and leading motors may dissociate from the filament and terminate the run. Interestingly, the run length of 2-motors attached to a flexible DNA scaffold (36 or 69 nm spacing) is significantly higher than when they are attached to the rigid DNA origami, suggesting that cargo rigidity may also play an important role in coordinating the myoVc motors. The velocity of 2- motor complexes were unchanged with motor spacing (Fig. 4C).

**FIGURE 4.**
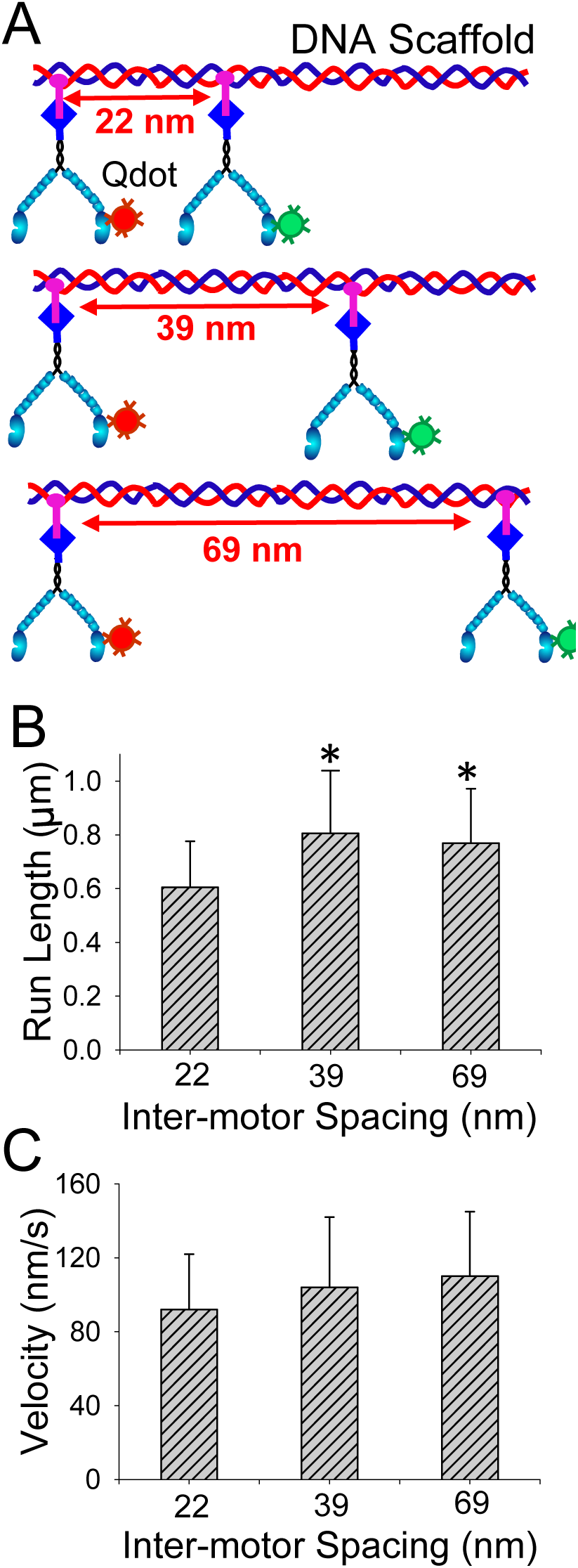
Effect of inter-motor spacing on two myoVc complexes. (A) Illustration of DNA scaffolds with 3 different inter-motor spacings (22, 39, or 69 nm). Motors with a C-terminal SNAP-tag were conjugated to a DNA scaffold containing SNAP-ligand. (B) Run lengths of the 2-motor complex on single actin filaments were significantly higher (*, p<0.05) when the spacing between motors was 39 nm (0.81±0.23 pm, N=68) or 69 nm (0.77±0.2 μm, N=72), compared with 22 nm (0.61±0.17, N=65) spacing. (C)Velocity was independent of inter-motor spacing: 22nm (92 ± 30nm/s, N=65), 39nm (104 ± 38nm/s, N=68) and 69 nm (110±35 nm/s, N=72). All values are reported ± SD.

### Two myoVc promote directional movement on actin bundles

The movement of a single myoVc versus a 2-myoVc complex was compared on actin bundles. The single myoVc had one motor domain labeled with a Qdot. For the 2-myoVc complex separated by 58 nm on a DNA origami, one motor domain of each myoVc was labeled with a different colored Qdot to allow unambiguous monitoring of the stepping dynamics of each motor (Fig. 1A). To examine the stepping dynamics at high temporal and spatial resolution, the Mg ATP concentration was reduced to 2μM to slow the stepping speed. A single myoVc was unable to move straight, switching between filaments in the bundle with almost every step (Fig. 5A). In addition to lateral steps, myoVc took a large number of back steps (26%) on the actin bundle (Fig. 5B,C). Steps of myoVc motors were identified by a step finding program (13), and step sizes were determined from the fit of a Gaussian distribution function. The mean forward step size of myoVc on the actin bundle was 63 ± 22 nm (mean ± SE, N=53) (Fig. 5C), similar to the step size of myoVa (63±26 nm, N=142)(13). The backward step size of myoVc on actin bundles (51± 20 nm, mean ± SE; N=19) was significantly shorter than that of the forward step size (Fig. 5C).

**FIGURE 5.**
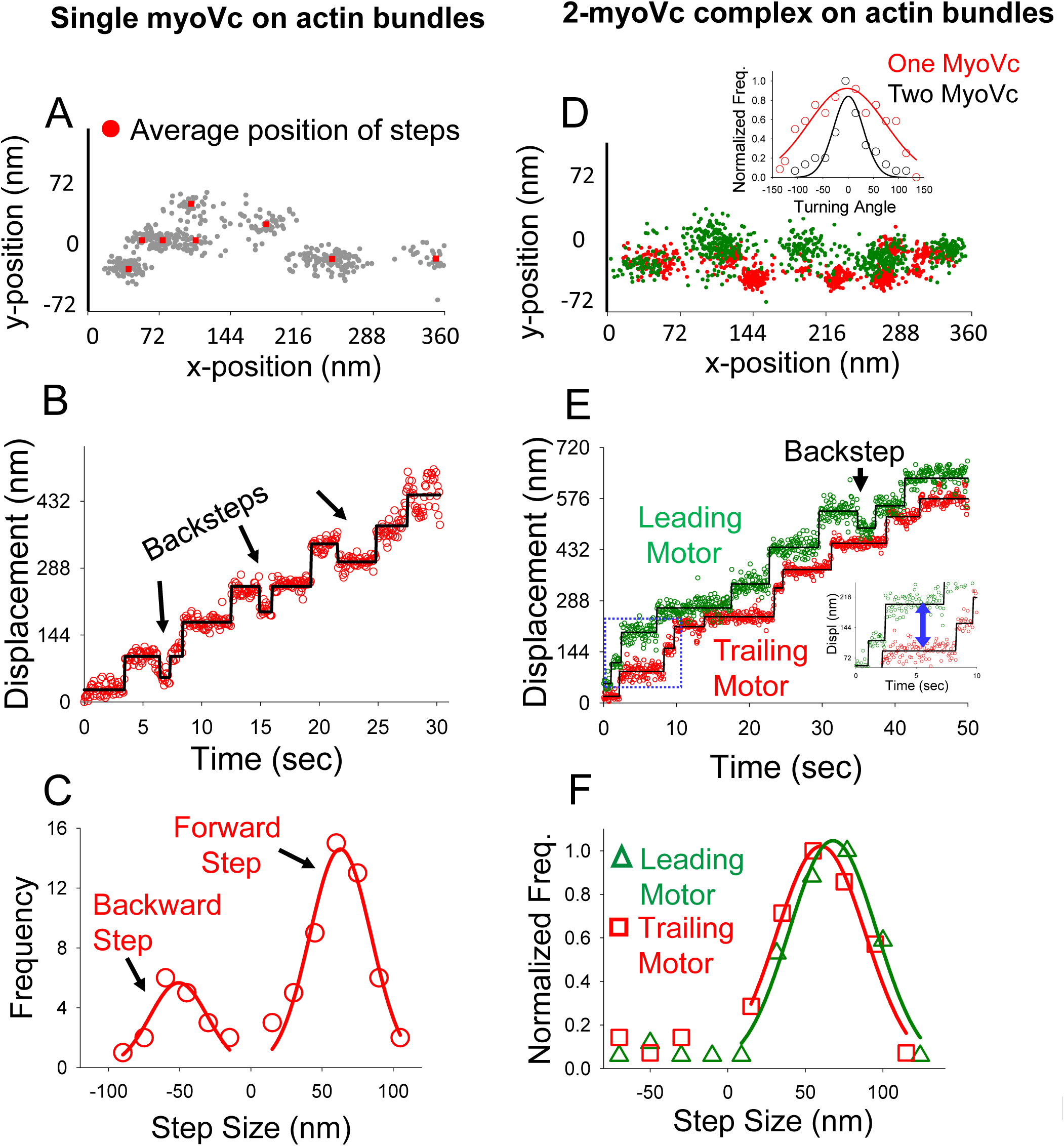
Motion of one versus two MyoVc motors on actin bundles at 2 μM MgATP. (A-C) Stepping dynamics of a single myoVc motor. (A) x,y trajectory (gray dots) of a single myoVc where the average position of each step is indicated by red dots. (B) A representative displacement versus time trajectory showed that a single MyoVc motor moves on actin bundles in a stepwise manner but with a significant number of back steps (arrow). (C) The mean forward 63±22 nm (mean±SE; N=53) and backward step sizes (51±20 nm, mean±SE; N=19) were determined from the fit of single Gaussian function. (D-F) Stepping dynamics of a 2-myoVc motor complex, in which one myosin is labeled with a red Qdot and the other with a green Qdot. (D) x,y trajectory of the leading (green) and trailing (red) motor within the 2-motor complex. (Inset) Histogram comparing the turning angles of a single motor (red) with the 2-motor complex (black). The average turning angles of a single and 2-motor complex were −2.4±76° (mean ± SD) and 0.31±29° (mean ± SD), respectively. (E) A representative trajectory showed that the leading (green) and trailing (red) motors within the complex moved on actin bundles in a stepwise manner. While stepping on the actin bundle, the inter-motor distance (blue arrow) fluctuates. The leading motor took an occasional back step (black arrow). (F) The forward and backward step-size distributions of the leading motor (green triangles). Forward and backward step-size distributions of the trailing motor (red squares).

Remarkably, the 2-myoVc complex moved much more directionally than a single motor (Fig. 5D). The directional movement of a single and 2- myoVc complex was quantified by measuring the turning angle between successive steps as done previously (2,15) (Fig. 5D, inset). The stepping pattern of a single myoVc motor showed that there is no bias in turning one direction over the other, determined from the x-y position (Fig.5A and D, inset). For single motors, 54% of the unbound head landed on the left side of the bound head, and the other 46% landed on the right side. Similarly, for the 2-myoVc complex, 53% of the unbound heads landed on the left side of the bound head, and 47% landed on the right side. The average turning angle of a single (-2.4 ± 76, mean ± SD, N=100) versus a 2-myoVc complex (0.31 ± 29, mean ± SD, N=64) was determined from the fit to a Gaussian distribution function. The smaller standard deviation (SD) of the 2-myoVc complex indicates that it moved more directionally than a single motor.

The step-sizes of both leading (68 ± 25 nm, mean ± SD, N=60) and trailing (60 ± 28 nm, mean ± SD, N=55) motors of the 2-myoVc complex were not statistically different than that of a single motor, but the frequency of the back steps was greatly reduced when 2-myoVc were physically linked (Fig. 5B). While the back step frequency of single motor was 26%, it was only ~8% for both leading and trailing motors within the 2-myoVc complex (Fig. 5F). The directional transport with reduced frequency of back steps or side steps suggests that myoVc motors cooperate to transport cargo along actin bundles.

### Forward and backward steps of myoVc on single actin filaments

To understand how multiple myoVc motors coordinate during cargo transport, the stepping dynamics of each Qdot-labeled motor within a 2-myoVc complex, coupled to a DNA origami with 58 nm inter-motor distance, was monitored on single actin filaments at low ionic strength (Fig. 6A). The mean step sizes of the leading (66 ± 27 nm, mean ± SD, N=95; Fig. 6B green) and trailing motors (61 ± 28 nm, mean ± SD, N=101; Fig. 6B red) within the complex were not significantly different (p<0.05) than a single motor stepping on an actin bundle (Fig. 6C). The mean backward step-size of both the leading (60 ± 19 nm, mean ± SD, N=11) and the trailing (56 ± 20 nm, mean ± SD, N=10) motors were slightly shorter than that of the forward steps (Fig. 6B). In addition to forward steps, both the leading and trailing motors take an almost same number of back steps(10% and 9%). This back step frequency is significantly lower than a single myoVc motor, and similar to the 2-myoVc complex moving on actin bundles (Fig. 5C, F). In contrast, when two myoVa motors were linked via a Qdot, the leading motor took a significantly higher number of back steps (11%) than the trailing motor (3%) (13).

**FIGURE 6.**
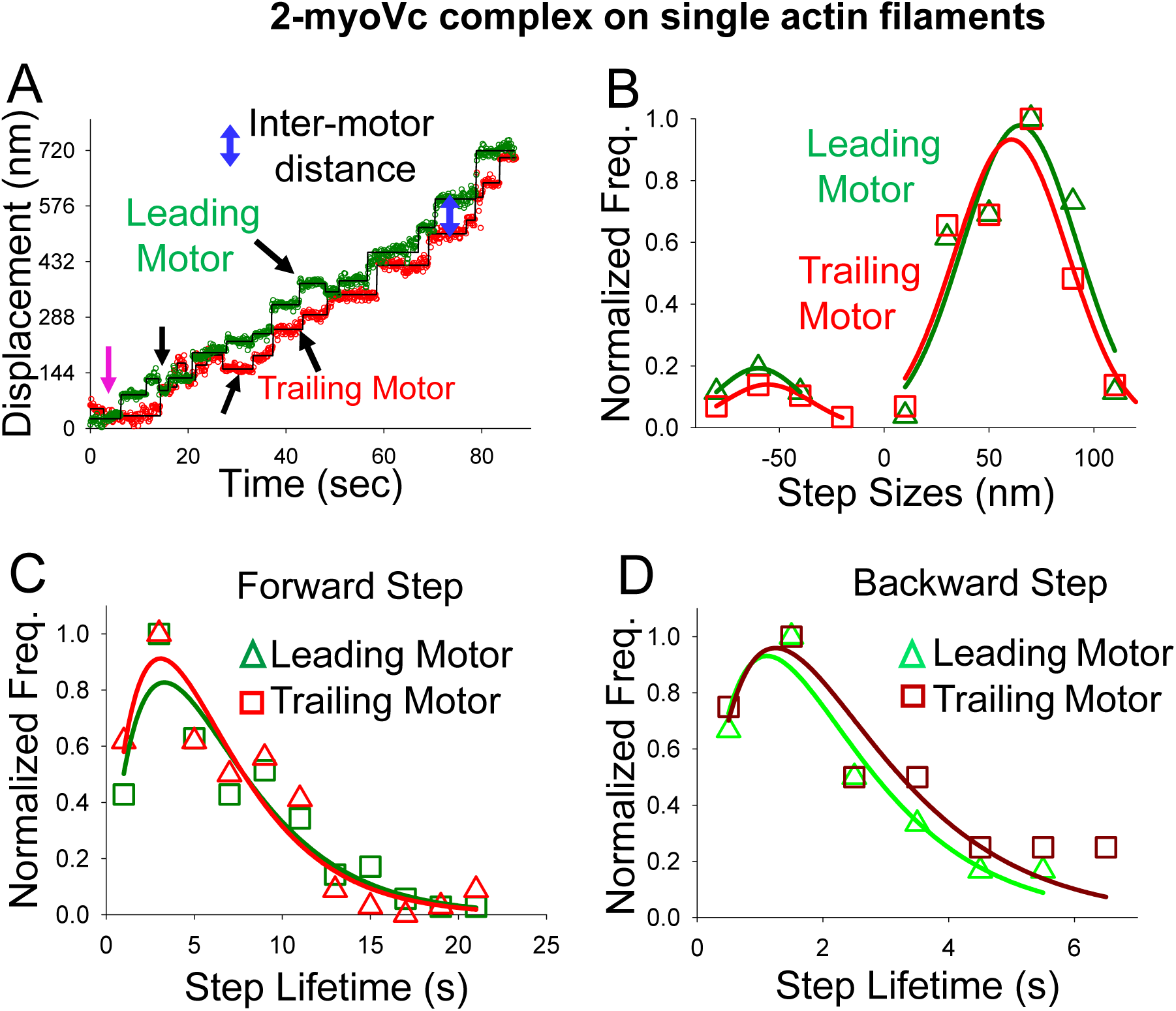
Characterization of a 2-myoVc complex stepping on single actin filaments. (A) A representative displacement versus time trajectory showed the stepwise movement of the leading (green) and trailing motor (red) of a two myoVc complex on an actin filament at 2 μM MgATP. While stepping, the distance between the two labeled heads fluctuates (blue double-headed arrow). (B) Step size distribution of the leading (green triangles) and the trailing motor (red squares) within a complex. The mean step sizes of the leading and the trailing motors were 66 ± 27 nm (N=95; green) and 61 ± 28 nm (N=101; red) respectively. Both leading and trailing motors take ~10% back steps. The forward (C) and backward (D) step lifetime distributions of the leading and trailing motors were the same.

The step lifetimes of the leading and trailing motors were plotted in histograms and fitted with P(t)= tk^2^e^-kt^ (see Experimental Procedures). Because both the leading and trailing motors are physically linked, the average step lifetime of both motors should be the same. As expected, the forward step lifetimes of the leading (3.3 ± 0.4s, mean ± SD, N=132) and the trailing (3.1 ± 0.5s, mean ± SD, N=133) motors are not significantly different (Fig. 6C). Similarly, the backward step lifetimes of both the leading and the trailing motors are the same (leading motor, 1.1 ± 0.2s, mean ± SD, N=15; trailing motor: 1.2 ± 0.3s, mean ± SD, N=14) (Fig. 6D).

### Inter-motor distance

While moving on an actin track, the distance between the two coupled myoVc motors fluctuates (Fig. 6A, 7A), despite being attached to the DNA origami with a 58 nm inter-motor spacing (16). The distance between two labeled heads of the leading and trailing motors was measured from the displacement versus time traces (Fig. 6A, blue arrow). As expected, the average distance between motors was 57 ± 51 nm (mean ± SD, N=277) (Fig. 7B). Due to the flexibility between the lever arm and rod domain, and the linkage between the origami and motor, the inter-motor distances fluctuated from −36 to ~ 160 nm (Fig. 7A, B). The negative inter-motor distance was occasionally seen when the trailing motor passed the leading motor at the beginning of the run (Fig. 6A, pink arrow), or when the trailing motor detached and reattached to the filament in front of the leading motor.

**FIGURE 7.**
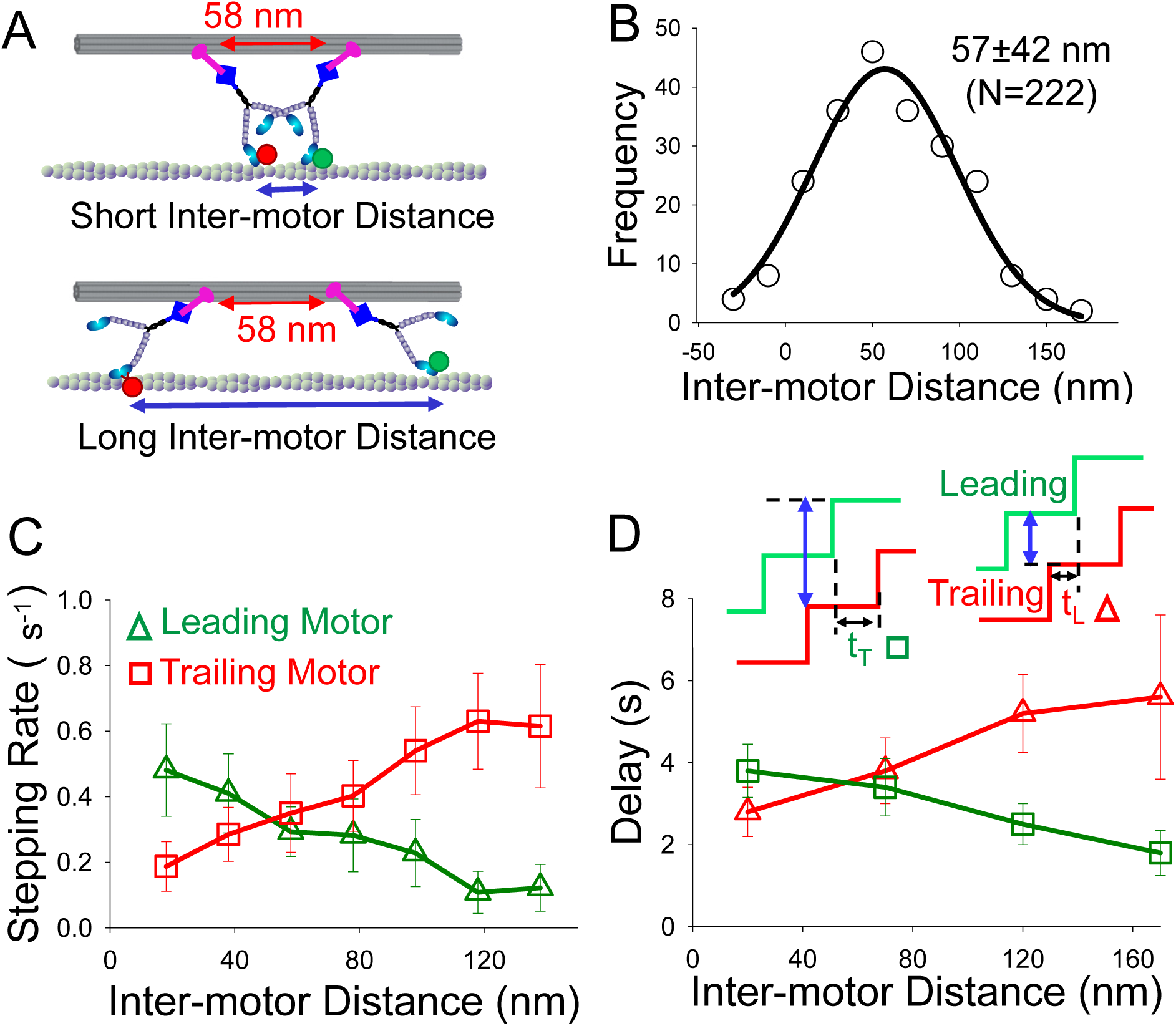
Characterization of inter-motor distances during myoVc stepping. (A) Illustration of a 2- motor complex showing the fluctuation of inter-motor distances on a single actin filament. (B) The distance between the labeled heads can fluctuate from −40 to 160nm, with a mean distance of 57 ± 42 nm (mean ± SD, N=222). (C) The stepping rate of both leading (green triangles) and trailing (red squares) motors gradually slow from the inter-motor distance 58nm to 78nm or 58nm to 38nm. At inter-motor distance >78nm, the stepping rate of the leading motor is slowed sharply, while the effect is opposite for the trailing motor. (D) Degree of coordination depends on the delay time between steps of the leading and the trailing motors. The average time delay (t_T_) between a forward step of the leading motor (green triangles) and the next forward step of the trailing motor (red squares) were plotted as a function of the inter-motor distance (blue arrow). These time delays were measured immediately after the step of the leading motor (green triangles). Similarly, time delay (t_L_) indicates the average time between a step of the trailing motor and the next step of the leading motor (red squares).

### Stepping rate and delay time between steps

To examine if myoVc motors coordinate their steps when mechanically coupled through a DNA origami, or if they move independently during transport (17), the stepping rate was measured as a function of inter-motor distance (Fig. 7C). The average stepping rate of the leading and trailing motor must be similar because both motors are physically linked. The stepping rate of both the leading and trailing motors are similar at intermotor distance 58±20 nm. (Fig. 7C). As the distance between motors increased further (>78 nm), the stepping rate of the leading motor decreased sharply, in contrast to the stepping rate of the trailing motor which increased. Surprisingly, the opposite effect was observed at short inter-motor distance (<38nm) where the leading motor’s stepping rate is higher than that of the trailing motor. This may be due to compression in the linkage between motors and/or a competition by the two motors for binding sites on the actin filament. A similar effect was observed when two myoVa motors were linked via a Qdot (13). This result suggests that as the inter-motor distance grows, leading and trailing motors tend to be pulled backward and forward from each other, respectively. Because the stepping rate changes as a function of inter-motor distance, the leading and trailing motors mechanically impact each other, and are not independent.

To further understand how the steps of the leading and trailing motors are coordinated, we measured the time delay (t_T_) for the trailing motor to take its step just after a step of the leading motor (13) (Fig. 7D). Interestingly, t_T_ is reduced as inter-motor distance is increased, suggesting that the trailing motor begins to coordinate its steps with that of the leading motor. The time delay for the leading motor to take a step just after the trailing motor (t_L_) increased with inter-motor distance, suggesting that the leading motor also coordinates with the trailing motor (Fig. 7D). These delay time and stepping rate relationships as a function of inter-motor distance suggest that the leading and trailing motors mechanically interact under tension.

## Discussion

Here we analyzed how the properties of two myoVc motors, coupled via a DNA origami as surrogate cargo, compared with the stepping of a single myoVc motor. Our results suggest that myoVc has unique attributes that specialize it for working in ensembles to move cargo on actin bundles. We first confirmed our previous finding that single molecules of myoVc can only move processively when on actin bundles, and provided that the ionic strength is low (2). Under no other conditions could we observe processive motion of a single myoVc. MyoVc surprisingly takes multiple back steps and lateral steps on bundles even in the absence of applied load. The stepping properties of processive myoVa differ in that back steps would only occur under load when it experiences a resistive force (18). We showed that the unbound head of myoVc switched filaments with almost every step when taking multiple steps on an actin bundle. The additional binding sites for the head in the bundle reduce the termination probability of the motor significantly. Additional binding sites are also useful for a single myoVa motor, which shows an increased run length on actin bundles compared with filaments (Fig. 2B) (15).

In contrast to the behavior of a single myoVc, coupling two myoVc motors via a common cargo produced continuous, unidirectional movement on an actin bundle. This result is similar to what Sakamoto and colleagues observed when a labeled scaffold with two coupled myoVc motors moved continuously on a single actin filament at low ionic strength (6). Our strategy for coupling the motors to the DNA origami furthermore allowed us to follow the stepping dynamics of one motor domain of each of the two coupled myoVc motors molecules, with high spatial and temporal resolution. Remarkably, the 2-myoVc complex essentially abolished the back steps and lateral steps taken by a single myoVc on an actin bundle. A potential mechanism is that because motors are mechanically coupled under tension, one motor can prevent the other from back steps or lateral steps. A similar mechanism was suggested for mammalian dynein. Whereas a single dynein motor in the absence of its activators dynactin and cargo adapters such as BicD (19) exhibits diffusional movement, coupling two dynein motors decreased the frequency of backward motion and favored directed transport along the microtubule (16). In cells, it is typical for multiple motors to work together to transport intracellular cargo, but in the case of myoVc multiple motor transport appears to be a necessity (17,20-22).

### Bundle preference

The preference of myoVc for actin bundles is evidenced by the observations that a single myoVc motor requires bundles to move processively at low ionic strength, and even small ensembles of myoVc require bundles to move continuously near physiologic ionic strength. Our previous work showed that elements in the lever arm/rod of myoVc were responsible for the large number of back steps that myoVc takes (2). The junction between the lever arm and rod of myoVc may be extremely flexible and cause an unfavorable geometry between the unbound head and the next binding site on a single actin filament, but allow forward motion given the additional lateral binding sites provided by an actin bundle. The step size of myoVc is also shorter (66 ± 27 nm, mean ± SD) than the helical pitch of the actin filament, in contrast to processive myoVa whose step size of 72 ± 12 nm on actin filaments more closely matches the helical pitch of the actin filament. Therefore, the lateral steps of myoVc may also result from the structural mismatch between the step size of myoVc and available actin binding sites. If the unbound head cannot bind to the next binding site on the same filament, it could bind to the adjacent filament within a bundle.

A single myoVc transitions from being non-processive on single filaments to processive on bundles, but the 2- and 4-myoVc complexes also show longer run lengths on bundles compared with filaments. This property may be an adaptation to help restrict the activity of myoVc to specialized actin cables, such as those assembled during the final stages of secretion in the exocrine pancreas (1). In contrast, 2- or 4-motor complexes of the myoVa isoform show shorter run lengths on bundles than on single actin filaments. It is possible that myoVa experiences an off-axis resistive load on an actin bundle which results in a higher termination rate from the track. MyoVa was shown to be sensitive when an off-axis resistive load was applied by an optical trap in a particular direction (23), but this property was never tested for myoVc.

### Run lengths of small ensembles

Run lengths of 2- and 4-myoVc complexes are 0.5 μm and 1.4 &3x03BC;m, whereas these values were much higher for myoVa (1.4 μm and 3.3 &3x03BC;m) on actin filaments. Why are the run lengths of two myoVc motors significantly lower than that of myoVa? A closer inspection of the stepping pattern of the 2-myoVa complex showed that each motor within the complex can detach and then reattach to the actin filament (13). Remarkably, a 2-motor complex of myoVa is driven by single motor for about ~50% of the time (13,14). When one of two myoVa motors detaches from the actin filament, the other attached motor is still capable of taking multiple steps on an actin filament. The detached motor can rebind and continue processive movement (13).

In contrast, we estimated that the 2-myoVc complex was driven by a single motor only a very small fraction of the time (~10%). Unlike myoVa, single molecules of myoVc cannot move processively on single actin filaments, and thus when one motor within the 2-myoVc complex detaches, the second motor simultaneously detaches. Therefore, it is expected that the run length of 2- or 4-motor complexes of myoVc will be significantly shorter than that of myoVa, as observed.

### MyoVc molecules coordinate to transport cargo

Our results showed that the velocity of the 2-myoVc complex was reduced significantly (28%), and the run length enhanced modestly, compared with a single myoVc, suggesting that the 2-motors are mechanically coupled under tension. Similar results were obtained when two myoVa motors were physically linked using a Qdot or a DNA scaffold (13,14). Our result is consistent with the observation that motors can negatively interfere with each other during motion (24). In contrast to these observations, it was suggested that if motors are independent of each other, the velocity will remain the same, the run length will increase several-fold, and the stall force will be a multiple of motor number (17,25,26). The mechanical interaction between two coupled motors also depends on the stiffness of the cargo and properties of the motor (14,27,28). A theoretical study suggested that the velocity, run length and detachment rate of motors depend on the internal strain between 2-motors and the stiffness of the cargo (20).

Because 2-motors are physically linked, it is expected that when the leading motor experiences a resistive load by the trailing motor, the trailing motor would also experience an assistive load by the leading motor. Interestingly, optical trapping studies showed that myoVa is more sensitive to resistive load than assistive load (29). This result is supported by the observation that the stepping rate of the leading motor decreased rapidly at higher inter-motor distance, whereas the stepping rate of the trailing motor increased. This suggests that while the leading motor experiences a resistive load by the trailing motor, the trailing motor experiences an assistive load by its partner motor. Thus, both the leading and trailing motors coordinate with each other and remain mechanically coupled under tension during cargo transport. Normally, myoVa takes back steps only when it experiences a resistive load. When 2- myoVa motors were linked via a Qdot, the leading myoVa took a higher number (11% versus 3%) of back steps than the trailing motor (13). In contrast, both the leading and trailing motors of myoVc took an equal number (~9%) of back steps, suggesting that due to the stiff linkage, both motors are exposed to the resistive load applied by its partner motor. Also, the step lifetimes of the leading and trailing motors of myoVc were not significantly different, although the backward stepping lifetime is 3 times shorter than that of the leading motor.

Interestingly, the delay time analysis with myoVc on actin filaments showed that both leading and trailing motors are affected by the inter-motor distance, suggesting that myoVc is equally sensitive to the resistive and assistive loads (Fig. 7D). In contrast, the delay time analysis of myoVa showed (13) that the trailing motor coordinate with the leading motor as the intermotor distance is increased, the leading motor does not shown to be coordinating with the trailing motor based on the delay time indicating that there is a differential effect on heads by the resistive load. We propose that when an ensemble of myoVc motors transport cargo, motors coordinate with each other and buildup strain through their linkage that impacts the stepping dynamics, and results a reduced velocity and longer run length.

### Conclusions

MyoVc is associated with secretory granules and likely involved in their trafficking (5). Transiently formed actin bundles, nucleated by formins at the plasma membrane, are the tracks on which zymogen granules are trafficked prior to secretion in the exocrine pancreas (1). This cellular observation is consistent with the findings here and those of a recent study (2), which showed that even teams of myoVc require actin bundles for continuous motion at physiologic ionic strength. The coordination we see with small ensembles of myoVc is thus likely to be a mechanism that ensures efficient secretory granule transport in the cell.

## Experimental Procedures

### Protein Expression, Purification, and Actin Bundle Preparation

Dimeric, constitutively active subfragments of human myoVc (amino acids 1-1105) and mouse myoVa (amino acids 1-1098) were cloned with a SNAP-tag at the C-terminus followed by a FLAG tag for affinity purification. Each heavy chain contained a biotin tag at the N-terminus for attachment of streptavidin Qdots. The biotin tag consisted of 88 residues from the *Escherichia coli* biotin carboxyl carrier protein (30). The heavy chain was co-expressed with a calcium-insensitive calmodulin (CamΔall) (31) to favor complete occupancy of the light chain binding sites, using the baculovirus/Sf9 cell system as described previously (2,32,33). Actin bundles were prepared using actin cross-linked protein fascin as described (15,32,34).

### Design of DNA origami nanotube

The DNA origami nanotubes were designed as previously described (11,16). In brief, one or two benzylguanine (SNAP-tag substrate) molecules per myoVa or myoVc binding site were attached to the staple strands corresponding to binding sites, i.e., columns 1, 13, 25, and 37 on the DNA origami. For imaging purpose, four Cy3 molecules were attached to the DNA which were located at one end (columns 1, 2, and 3) of the DNA origami. The folding was conducted by mixing the ‘base staple mix’ containing Cy3-labeled staple strands, the benzylguanine staples and viral singlestranded DNA (M13mp18 DNA) in TAE Mg^2^+ buffer (40 mM Tris-acetate pH 8.0, 1 mM EDTA and 4mM Mg(OAc)_2_). Then, the mixture was heated at 80°C for 5 min, followed by two-step cooling from 80°C to 60°C. The DNA origami contains 1, 2, or 4 binding sites that are separated by 58 nm. DNA scaffolds were prepared as described (6).

### Conjugation of motor proteins with a DNA origami nanotube or DNA scaffold

Before conjugation with motor proteins, DNA origami nanotubes were purified with a centrifugal filter device. Briefly, 100 μl of DNA origami nanotubes were transferred to an Amicon Ultra-0.5 filter device (Millipore, 100 kDa MWCO) and washed four times with TAE Mg^2+^ buffer according to the manufacturer’s instructions. The concentration of DNA was measured spectrophotometrically at 260 nm. DNA was precipitated by adding 8%(w/v) PEG8000 (MP Biomedicals) in buffer A (25 mM imidazole, pH 7.4, 4 mM MgCl_2_, 1 mM EGTA, 25 mM KCl, 10 mM DTT) and spun at 15,000g for 15 min at 22°C. The precipitated DNA was resuspended in buffer A, mixed with myoVc (or myoVa) at a molar ratio of 5:1 (protein to binding sites), and incubated for 1 h at 27°C. The unbound motor proteins were removed by passing through ~450 μl of Sephacryl S-500 HR (GE Healthcare) packed in a spin column at 1,000 x g for 12 s. Conjugation of SNAP-tagged myoVc or myoVa to a double-stranded DNA scaffold was carried out as described (35). The DNA origami-motor complex was diluted to 1 μM of myoVc or myoVa motor for experiments.

All DNA origami nanotubes were labeled with Cy3 dyes which were attached to one end of the nanotube. DNA origami containing a single motor or 4 motors was detected from Cy3 fluorescence using TIRF microscopy. For preparing the 2-motor complex (Fig. 1A), we mixed 2 pl of 1 pM streptavidin conjugated red (emits at 655 nm) and 2 μl of 1 μM green Qdots (emits at 655 nm), into 1 μl of 1 μM myoVa or myoVc (origami-motor complex) in buffer B (buffer A plus mg/ml BSA), and incubated for 15 min at room temperature. Streptavidin conjugated Qdots were attached specifically to the biotinylated head domain of the motor. A 4-fold molar excess of Qdot over motor was used so that all motors would be labeled with Qdots. If both binding sites on the DNA origami are occupied by motors labeled with Qdots, there will be four equal probabilities: red:red, green:green, red:green, and green:red. Therefore, it is expected that a maximum of 50% of the DNA origamis will be dual color (red and green). We counted the number of Qdot labeled motors bound to actin filaments and showed that ~40% (N=165) of complexes were labeled with dual color Qdots (Fig. 1C). This analysis suggests that the binding affinity of myoVc motors to the origami and labeling with Qdots were both very efficient (80%). The experiments of the 2-motor complex were conducted either at 1 mM MgATP for run length and velocity measurements, or at 2 μM MgATP for resolving the individual motor steps.

### Microscopy and Data Analysis

*In vitro* motility experiments were performed at room temperature (23 ± 1°C) using a Total Internal Reflectance Fluorescence (TIRF) microscope as described previously (36). Experiments were performed in buffer A (25 mM imidazole, pH 7.4, 4 mM MgCl_2_, 1 mM EGTA, 25 mM KCl, 10 mM DTT) except where noted. Qdots and actin filaments were excited with a 488 nm argon laser. Typically, 1000 images were captured at 10-30 frames/second (1 pixel = 117 nm) using an intensified CCD camera (Stanford Photonics, CA). Qdots were tracked using the SpotTracker 2D or MtrackJ plugins of Image J 1.41v to generate motion paths in two dimensions or to measure the total run length (National Institutes of Health, Bethesda, MD) as described (36). Digital images (red and green Qdots) were corrected for color registration error as described (36). Quantum dot labeled motors were tracked with high spatial (6 nm) and temporal (33 ms) resolution as described previously (32).

Velocity and run length of Qdot labeled motors were calculated as described previously (36). For step size measurements, experiments were performed at low ATP concentration (2 μM MgATP). Qdot labeled motors were tracked using the SpotTracker 2D which provides x,y data of the motion. We then plotted displacement versus time trajectories. Using a step-finding program (37), the step sizes and step lifetime were determined. On actin bundles, step sizes were measured from the average x,y values which were identified using step finding program from the time versus displacement trajectory. Step sizes were plotted in a histogram and fitted to a Gaussian distribution function. The step lifetime of both leading and trailing motors was plotted in a histogram and fitted using the equation P(t)= tk^2^e^-kt^, where 1/k is the step lifetime (38). For statistical significance, Student’s t-test for velocity, and Kolmogorov–Smirnov test for run length and step lifetime comparisons were used. Stepping rate and delay time of motors were measured as a function inter-motor distance as described (13).

## ACKNOWLEDGEMENTS

We thank David Warshaw for the use of the TIRF microscope, Andrej Vilfan for insightful comments and Guy Kennedy for technical assistance. This work was supported by the American Heart Association (12SDG11930002, MYA), and Grant-in-Aid for Scientific Research, Japan (15KT0155, KF) and the National Institutes of Health (GM078097, KMT).

## AUTHOR CONTRIBUTIONS

M.Y.A. and K.M.T. designed research; M.Y.A., E.B.K. and K.F. performed experiments; M.Y.A., K.M.T., K. F. and K. O. contributed analytical tools; M.Y.A. analyzed data. M.Y.A., K.M.T. and K. F. wrote the article. E.B.K. and K.F. contributed equally to this work. All authors approved the final version of the manuscript.

